# Cross-sectional and longitudinal associations of threat and deprivation on cognition, emotional processing and psychopathology in children and adolescents

**DOI:** 10.1101/2020.02.09.940858

**Authors:** Julia Luiza Schäfer, Katie A. McLaughlin, Gisele Gus Manfro, Pedro Pan, Luis Augusto Rohde, Eurípedes Constantino Miguel, Giovanni Abrahão Salum

## Abstract

**Background:** Exposure to childhood adversity has been consistently associated with poor developmental outcomes, but it is unclear whether these associations vary across different forms of adversity. We examined cross-sectional and longitudinal associations between two types of adversity—threat and deprivation—with cognition, emotional processing, and psychopathology in a middle-income country.

**Methods:** The sample consisted of 2,511 children and adolescents (6-17 years old) from the Brazilian High-Risk Cohort for Psychiatric Disorders. Parent reports on childhood adversity were used to construct threat and deprivation latent constructs. Psychopathology was measured by the CBCL which generated a measure of general psychopathology (the “p” factor). Executive function (EF) and attention orienting toward angry faces were assessed using cognitive tasks. All measures were acquired at two time-points 3-years apart. Cross-lagged panel models were estimated to evaluate longitudinal associations.

**Results:** Psychopathology was associated with threat and deprivation cross-sectionally, and higher levels of threat and deprivation predicted increases in general psychopathology at follow-up. For EF, worse performance was more strongly associated with deprivation than threat at baseline, and only with deprivation at follow-up. Deprivation was associated with attention orienting away from angry faces cross-sectionally, but neither form of adversity was associated with changes over time in attention bias.

**Conclusion:** Both types of adversity are associated with current and future psychopathology and with current, but not future, EF. Threat was more strongly linked to higher psychopathology, whereas deprivation was more strongly linked to lower EF. Lack of longitudinal associations between deprivation and EF mean reverse causality cannot be excluded.

## Introduction

Childhood adversity involves negative environmental experiences that require considerable adaptation by an average child, including physical, emotional, and sexual abuse, neglect, domestic violence, and parental absence (McLaughlin, 2016). These experiences are highly prevalent around the world (Costello *et al.*, 2002) and especially in low and middle income countries (Viola *et al.*, 2016). Exposure to childhood adversity represents a public health problem due to its extensive costs to the society and individuals (Magruder, McLaughlin and Elmore Borbon, 2017), leading to poorer mental health (Green *et al.*, 2010; McLaughlin et al., 2012) and academic achievement, such as lower grades, higher school-days absence, and more frequent suspensions (Lansford *et al.*, 2002). Determining how distinct adverse childhood experiences influence emotional and cognitive development is critical to identify novel strategies for preventing the emergence of psychopathology and developmental problems in children who have experienced adversity.

Previous findings have suggested that different types of adverse experiences might influence development in different ways and, therefore, should be distinguished (Mclaughlin and Sheridan, 2016). One prominent model proposes that childhood adversities encompass two types of abnormal inputs (Humphreys and Zeanah, 2014): *threat*, the presence of an unexpected input that represents threat to the physical integrity, or well-being of the child; and *deprivation*, the absence of some type of expected input (McLaughlin, Sheridan and Lambert, 2014; McLaughlin, 2016). Examples of experiences involving high levels of threat are physical, sexual, and emotional abuse, witnessing domestic violence, and exposure to violence in the community, or at school; while experiences involving low levels of cognitive and social stimulation, such as neglect, institutional rearing, other forms of parental absence, and material deprivation associated with chronic poverty are characterized by high levels of deprivation (McLaughlin, Sheridan and Lambert, 2014; Sheridan and Mclaughlin, 2014; McLaughlin, 2016; McLaughlin *et al.*, 2019).

A growing number of studies have showed that threat and deprivation influence child development differently. More specifically, children who have experienced violence, and not deprivation, require less perceptual information to identify anger (Pollak *et al.*, 2000; Pollak and Sinha, 2002), classify a wider range of negative emotion as anger (Pollak and Kistler, 2002), and exhibit attention biases to threatening social information (Shackman and Pollak, 2014). On the other hand, exposure to deprived environments has been associated with poor performance on complex cognitive tasks, particularly those encompassing language abilities and executive function (EF) (Mclaughlin, Sheridan and Nelson, 2017; Lawson, Hook and Farah, 2018). Hence, while threat seems to have strong influences on emotional processing—particularly in relation to cues that are potentially threatening, deprivation seems to be associated with reduced performance on complex tasks reflecting EF (McLaughlin, 2016). Moreover, as it is of knowledge, the relationship between childhood adversity and psychopathology is already well-established. Although the dimensional model does not make differential predictions about how threat and deprivation influence psychopathology, emerging evidence from population-based longitudinal studies, suggests that threat may have stronger associations with psychopathology than deprivation (Platt *et al.*, 2018).

The previous literature is limited in two important ways. First, most research using this theoretical framework has focused on youth living in high-income countries. However, most of the world’s youth live in poor countries and face adversities for which there is limited data available (Viola *et al.*, 2016). Second, most existing research investigating the associations between childhood adversity, psychopathology, and cognition is cross-sectional and does not examine how these experiences influence developmental trajectories of emotion, cognition, and psychopathology or address the possibility of bias due to reverse causality.

In this study we examined the longitudinal associations of threat and deprivation with cognition, emotion, and psychopathology in children and adolescents in a large school-based community sample from a middle-income country. Specifically, we aimed to: (1) to investigate the goodness of fit of the threat and deprivation model in a large community sample from Brazil; and (2) to investigate the associations of threat and deprivation with EF, emotional processing as measured by attention orienting toward angry faces, and psychopathology. We hypothesized that a model specifying distinctions among adversities reflecting threat and deprivation would provide a good fit to the data. We also expected that attention bias toward angry faces would be associated with threat, but not with deprivation; that worse EF would be associated with deprivation, but not with threat (McLaughlin, 2016), and that psychopathology would be associated with both threat and deprivation

## Methods and Materials

### Study design, procedures and participants

Data for this study is drawn from the baseline and 3-year follow-up waves of the Brazilian High-Risk Cohort for Psychiatry Disorders (BHRCS), a school-based community cohort from the cities of São Paulo and Porto Alegre. Briefly, in the year 2010, 9937 parents of 6-14-years-old children from 57 schools in São Paulo and Porto Alegre were screened using the Family History Survey (FHS; Weissman *et al.*, 2000). From this sample, two subgroups were recruited for further assessments. One subgroup was randomly selected (n=957), while the other was selected from a high-risk score procedure used to identify children with current symptoms and/or family history of psychiatric disorders (n=1554). A total of 2,511 children/adolescents and their parents were assessed at two time points through questionnaires and interviews about the history of exposure to adversities, and psychopathology. Children/adolescents also completed neurocognitive tests at both time points. The Institutional Review Boards of all institutions involved in the study have given approval for the research protocol, and every participant, parents, or caregivers who participated has given written, or verbal informed consent. For a detailed description of the study, its procedures, and sample see Salum *et al.*, 2014.

### Measures

#### Exposure to Adversity

Selected variables from the BHRCS were chosen based on theoretical models of adversity. A detailed description of the selected questions and response options is depicted in the Supplemental Table S1. All baseline data was drawn considering lifetime exposure, while follow-up data was drawn considering, only, exposure over the last three years from baseline to the follow-up assessment.

Variables selected for measuring exposure to threat were drawn from two parent report sources. First, exposure to physical and sexual abuse, attack or threat, witnessing domestic violence and witnessing attack, were investigated using the Posttraumatic Stress Disorder (PTSD) assessment of the Development and Well-Being Assessment (DAWBA; Goodman *et al.*, 2000). Second, questionnaires specifically designed for the BHRCS were used to evaluate exposure to bullying, and the frequency of exposure to psychological, physical and sexual abuse (never, once or twice, from time to time, and often) (Salum *et al.*, 2016).

Exposure to deprivation experiences was measured through the assessment of the mother’s educational level (ranging from without any study to postgraduate education), family income (measured in quintiles), socioeconomic classification (A/B – the wealthiest, C, or D/E – the poorest), father contact (in contact, non-contact, deceased, or unknown), and the frequency (never, once or twice, from time to time, and often) of exposure to neglect (Salum *et al.*, 2016).

### Cognition

#### Executive Function

Three dimensions of executive function (EF), measured by two tasks each, were calculated in order to create a second order model of EF. The dependent variable was a single EF standardized score encompassed by working memory, inhibitory control, and temporal processing scores after regressing out effects of age and gender on the task parameters. Higher scores represent better EF.

*Working memory* was measured by the *digit span* (a subtest of the WISC-III; Wechsler, 2002) and *corsi blocks* tasks (Vandierendonck *et al.*, 2004). Both tasks involve the repetition of a given sequence. While in the *digit span task* the participants hear and repeat an increasingly difficult sequence of numbers, either forward and backward, in the *corsi blocks task* they repeat an increasingly difficult spatial sequence tapped by a researcher on up to nine identical blocks. Both outcomes are the level at which a correct repetition failed twice consecutively.

*Inhibitory control* was measured by the *conflict control task* (CCT; Hogan *et al.*, 2005) and the *go/no-go* task (GNG; Bitsakou *et al.*, 2008). Both consist of an arrow based visual stimuli with a total 100 trials divided into two different instructions. In the *conflict control task*, participants are asked to press a button indicating the direction or opposite direction of arrows shown on the screen. Participants either press the button indicating the correct direction of a green arrow (75 congruent trials), or press the button indicating the opposite direction of a red arrow (25 incongruent trials). The *go/no-go* task requires participants to completely suppress the tendency to press the buttons indicating the direction of the green arrows (75 *go* stimuli trials) when a double-headed green arrow (25 *no-go* stimuli trials) appears on the screen. For both tasks, intertrial interval was 1,500 ms, and the stimulus duration was 100 ms. The outcomes were the percentage of correct responses in the incongruent trials (CCT) and the percentage of failed inhibitions in the *no-go* trials (GNG).

Finally, *temporal processing* was measured by both *time anticipation (TA) tasks 400 ms and 2,000 ms* (Toplak and Tannock, 2005) on baseline, and only *time anticipation* 400 *ms* on the follow-up. These tasks require participants to anticipate when a visual stimulus will appear. In a game-like manner, the task involves an allied spaceship running out of oxygen and the participant has to give it to them in order to save the crew. In each task, the allied spaceship is visible for the first 10 trials, while for the remaining 16 trials the spaceship is invisible due to an invisible shield. Then, participants are asked to press a button to anticipate when it arrives. A 750-ms window of time to respond correctly and feedback after every trial are given. In task 1, the anticipation interval is 400ms, while in task 2 it is 2,000ms. The outcome is the mean percentage of button pressed in the correct time window interval for the invisible part of the task. Tasks involving temporal delays with flexible cognitive demands have been proposed to be a part of EF in some models (Barkley, 1997). Temporal processing tasks used have previously been well-correlated with the other EF tasks in our sample (Martel *et al.*, 2016; Manfro *et al.*, 2019). Results for EF model fit are reported in the Supplemental Material.

### Emotional Processing

#### Attention orienting toward angry faces

Attention orienting toward angry faces was assessed using a dot-probe task in Eprime 2.0 (Psychology Software Tools, USA). The task consists of two versions, one longer than the other, of the same identical stimuli presentation, which are paired threatening (angry) and neutral face photographs followed by a probe at the location of one of the two photographs. In both versions, each trial starts with a central fixation cross (for 500ms), followed by the face pair (for 500 or 1250 ms) which is replaced with the probe (for 1100ms). Participants are instructed to press one of the response keys to indicate whether the probe appeared in the left, or right side of the screen. Trials are, randomly, either congruent (16 trials for the short version and 32 for the long version), with threatening faces and probes appearing on the same side of the screen, and incongruent (16 trials for the short version and 32 for the long version), with threatening faces appearing on opposite sides of the screen. The inter-trial interval varies randomly from 750 to 1250ms. Since the neutral and the threatening stimuli are in different screen locations, they compete for attention. Therefore, attention orienting toward angry faces is measured as the difference in reaction time between the task’s trials in which the probe replaces a neutral stimulus versus those in which the probe replaces a threatening stimulus. Response times were excluded as errors from trials where the response was incorrect or did not occur before probe offset. Additionally, response times less than 200ms or more than 2 standard deviation above each participant’s mean were excluded as outliers, as well as attention bias scores were not calculated if more than 50% response times data were missing. Therefore, the dependent variable was a standardized score of attention orienting toward angry faces after regressing out effects of age and gender on the task parameters. Scores greater than zero represent biases in attention toward threats and lower than zero biases in attention away from threats.

### Psychopathology

Psychopathology was measured dimensionally at baseline and follow-up through the Child Behavior Checklist (CBCL; Achenbach and Rescola, 2001; Bordin *et al.*, 2013). The CBCL is a parent-report questionnaire that assesses the child’s emotional, behavioral, and social problems yielding a total score (including all items), as well as an internalizing and externalizing score. A bifactor model with one dimension of general psychopathology (the “p” factor) was fitted to the data with two residualized dimensions of internalizing and externalizing psychopathology. Only general psychopathology scores were used for further analysis. More detail on the model’s structure are described in the Supplemental Material.

## Data Analysis

First, we conducted factor analyses in order to assess the latent structure of threat and deprivation models of exposure to adversity, as well as the EF and psychopathology models. Missing data was accounted for using full information maximum likelihood estimation. Model goodness of fit was evaluated using root-mean-square error of approximation (RMSEA), comparative fit index (CFI), and Tucker-Lewis index (TLI). According to the literature, a RMSEA equal or below .06, and a CFI and a TLI above .95 indicate good fit (Hu and Bentler, 1999; Kline, 2013). Second, we used the observed factor scores from the validated models to test the longitudinal associations of threat and deprivation with psychopathology, cognition, and emotional processing through Cross-Lagged Panel Models (CLPM) with baseline and 3-year follow-up assessments. Given that EF was comprised of multiple distinct tasks, we additionally examined associations of threat and deprivation with each of the three components of EF. Such analysis was chosen to investigate longitudinal associations due to CLPM estimation of the directional influence variables have over time, while controlling for correlations within time-points and autoregressive effects (or stability) across time, which provide a robust strategy for estimating longitudinal associations and minimizing bias due to reverse causality (Kearney, 2018). Data analysis were performed using the *Mplus* software for categorical data and the *lavaan* package from R for continuous data and CLPMs.

## Results

### Descriptive Statistics

The total sample was comprised of 2,511 children and adolescents with a mean age of 10.42 years old at baseline and 13.71 years old on follow-up. Among those, 1375 (54.8%) were female, and 1256 (50.1%) were from the city of São Paulo. Descriptive data on variables of interest are shown on Table 1 and additional descriptive data is found in the Supplemental Material.

**Table 1.** Sample description.

|  | Baseline |  |  | 3-year follow-up |  |  |  |  |
| --- | --- | --- | --- | --- | --- | --- | --- | --- |
|  | N | % | Valid N |  |  |  |  |  |
| Sex: male | 1375 | 54.8 | 2511 |  |  |  |  |  |
| Site: São Paulo | 1256 | 50 | 2511 |  |  |  |  |  |
|  | Mean (sd) | Min | Max | Valid N | Mean (sd) | Min | Max | Valid N |
| Age (years) | 10.2 (1.9) | 5.83 | 14.37 | 2511 | 13.5 (1.9) | 9.2 | 17.87 | 2010 |
| Family income (BRL) | 757.6 (536.6) | 29.94 | 4910.18 | 2110 | 783.8 (656.8) | 21.65 | 8658.01 | 1679 |
| CBCL: Total CBCL scores | 17.2 (16.1) | 0 | 101 | 2511 | 14.6 (15.0) | 0 | 90 | 2010 |
| Working Memory |  |  |  |  |  |  |  |  |
| Corsi block (backward) | 4.8 (2.1) | 0 | 14 | 2223 | 5.6 (2.5) | 0 | 13 | 1880 |
| Digit span (backward) | 3.5 (1.6) | 0 | 12 | 2249 | 4.1 (2.0) | 0 | 13 | 1880 |
| Inhibitory Control |  |  |  |  |  |  |  |  |
| CCT % Correct Inhibitions | 0.6 (0.2) | 0 | 1 | 2165 | 0.7 (0.3) | 0 | 1 | 1704 |
| Go/No-Go: Comission | 0.3 (0.2) | 0 | 1 | 2158 | 0.2 (0.2) | 0 | 1 | 1701 |
| Temporal Processing |  |  |  |  |  |  |  |  |
| Time Anticipaion (0.4s): hits | 0.6 (0.2) | 0 | 1 | 2185 | 0.8 (0.2) | 0 | 1 | 1701 |
| Time Anticipaion (2s): hits | 0.3 (0.2) | 0 | 1 | 2180 | NA | NA | NA | NA |
| Attention orienting toward angry faces (ms) | 5.2 (53.0) |  |  | 2148 | 2.9 (40.0) |  |  | 1603 |

**Note:** crude scores for psychopathology, executive function tasks, and attention orienting toward angry faces are presented in order to inform about the variables' characteristics on the sample.

### Threat and Deprivation Latent Structure

The model of threat and deprivation as latent variables (Supplemental Figure S1) was tested at both the baseline and follow-up assessments. The baseline model, consisting of nine variables informing the threat dimension and five variables informing the deprivation dimension, showed acceptable fit indexes (CFI = 0.942, TLI = 0.928, RMSEA = 0.036). For the follow-up model, frequency of exposure was considered for the three-year time frame between baseline and follow-up assessments. Due to low frequency on both variables assessing sexual abuse exposure over the last three years, these variables were combined. Therefore, the follow-up model consisted of eight variables informing the threat dimension and five variables informing the deprivation dimension, presenting acceptable goodness of fit (CFI = 0.923, TLI = 0.903, RMSEA = 0.037). Detailed information is provided on Table 2.

**Table 2.** Factor loadings of the Threat and Deprivation Models at baseline and follow-up.

|  | Baseline |  |  |  | Follow-up |  |  |  |
| --- | --- | --- | --- | --- | --- | --- | --- | --- |
|  | Estimate | S.E. | Est./S.E. | *p*-value | Estimate | S.E. | Est./S.E. | *p*-value |
| **Threat** |  |  |  |  |  |  |  |  |
| Bullying exposure | 0.458 | 0.033 | 13.687 | <0.001 | 0.436 | 0.043 | 10.090 | <0.001 |
| DAWBA: Physical abuse | 0.838 | 0.035 | 23.741 | <0.001 | 0.848 | 0.042 | 19.987 | <0.001 |
| Physical abuse | 0.654 | 0.035 | 18.630 | <0.001 | 0.634 | 0.047 | 13.539 | <0.001 |
| Emotional abuse | 0.532 | 0.029 | 18.263 | <0.001 | 0.619 | 0.036 | 17.207 | <0.001 |
| DAWBA: Sexual abuse | 0.555 | 0.079 | 7.044 | <0.001 | - | - | - | - |
| Sexual abuse | 0.576 | 0.062 | 9.293 | <0.001 | - | - | - | - |
| Sexual abuse (combined) | - | - | - | - | 0.363 | 0.079 | 4.624 | <0.001 |
| DAWBA: Attack or threat | 0.532 | 0.049 | 10.803 | <0.001 | 0.544 | 0.054 | 10.106 | <0.001 |
| DAWBA: Domestic violence witnessing | 0.682 | 0.037 | 18.452 | <0.001 | 0.736 | 0.043 | 17.038 | <0.001 |
| DAWBA: Attack witnessing | 0.690 | 0.042 | 16.300 | <0.001 | 0.740 | 0.049 | 15.002 | <0.001 |
| **Deprivation** |  |  |  |  |  |  |  |  |
| Mother's educational level | 0.355 | 0.030 | 11.835 | <0.001 | 0.411 | 0.058 | 7.072 | <0.001 |
| ABEP 2009: Stratified Score | 0.641 | 0.035 | 18.129 | <0.001 | 0.576 | 0.077 | 7.501 | <0.001 |
| Father status | 0.406 | 0.041 | 9.821 | <0.001 | 0.235 | 0.060 | 3.881 | <0.001 |
| Neglect | 0.621 | 0.038 | 16.148 | <0.001 | 0.465 | 0.071 | 6.527 | <0.001 |
| Family income | 0.756 | 0.039 | 19.427 | <0.001 | 0.842 | 0.108 | 7.775 | <0.001 |
| **Model fit baseline:** CFI= 0.942, TLI= 0.928, RMSEA= 0.036; **Model fit follow-up:** CFI= 0.923, TLI= 0.903, RMSEA= 0.037 |  |  |  |  |  |  |  |  |

### Cross-sectional and cross-lagged effects from the Cross-Lagged Panel Models

The analysis investigating effects of threat and deprivation and psychopathology, EF and attention orienting toward angry faces from the CLMP are depicted in Table 3 and summarized below.

**Table 3.** Cross-lagged Panel Models results.

| Cross-sectional |  |  |  |  |  |  |
| --- | --- | --- | --- | --- | --- | --- |
|  | Baseline |  |  | Follow-up |  |  |
| | $\beta$ | *p value* | CI 95% | $\beta$ | *p value* | CI 95% |
| Threat |  |  |  |  |  |  |
| Psychopathology | 0.430 | <0.001 | 0.398, 0.462 | 0.349 | <0.001 | 0.311, 0.388 |
| Executive Function | -0.086 | <0.001 | -0.125, -0.046 | 0.014 | 0.532 | -0.031, 0.060 |
| Attention Bias | 0.033 | 0.125 | -0.009, 0.075 | 0.042 | 0.092 | -0.007, 0.091 |
| Deprivation |  |  |  |  |  |  |
| Psychopathology | 0.160 | <0.001 | 0.122, 0.198 | 0.058 | 0.010 | 0.014, 0.101 |
| Executive Function | -0.155 | <0.001 | -0.194, -0.116 | -0.061 | 0.008 | -0.106, -0.016 |
| Attention Bias | -0.027 | 0.210 | -0.069, 0.015 | -0.057 | 0.024 | -0.106, -0.008 |
| Longitudinal associations |  |  |  |  |  |  |
|  | Adversity → Outcome |  |  | Outcome → Adversity |  |  |
| | $\beta$ | *p value* | CI 95% | $\beta$ | *p value* | CI 95% |
| Psychopathology |  |  |  |  |  |  |
| Threat | 0.121 | <0.001 | 0.076, 0.167 | 0.123 | <0.001 | 0.078, 0.168 |
| Deprivation | 0.064 | 0.003 | 0.022, 0.107 | 0.038 | 0.047 | 0.000, 0.076 |
| Executive Function |  |  |  |  |  |  |
| Threat | -0.031 | 0.160 | -0.074, 0.012 | -0.031 | 0.150 | -0.074, 0.012 |
| Deprivation | -0.041 | 0.069 | -0.085, 0.003 | -0.056 | 0.002 | -0.091, -0.021 |
| Attention bias |  |  |  |  |  |  |
| Threat | -0.044 | 0.096 | -0.096, 0.008 | -0.010 | 0.650 | -0.055, 0.034 |
| Deprivation | 0.013 | 0.619 | -0.039, 0.066 | -0.030 | 0.109 | -0.067, 0.007 |
| Autoregressive |  |  |  |  |  |  |
|  | Autoregressive |  |  |  |  |  |
| | $\beta$ | *p value* | CC 95% | | | |
| Psychopathology | 0.336 | <0.001 | 0.295, 0.378 |  |  |  |
| Threat | 0.280 | <0.001 | 0.234, 0.326 |  |  |  |
| Deprivation | 0.638 | <0.001 | 0.609, 0.667 |  |  |  |
| Executive Function | 0.435 | <0.001 | 0.398, 0.473 |  |  |  |
| Threat | 0.330 | <0.001 | 0.288, 0.371 |  |  |  |
| Deprivation | 0.631 | <0.001 | 0.601, 0.660 |  |  |  |
| Attention bias | 0.064 | 0.025 | 0.008, 0.120 |  |  |  |
| Threat | 0.331 | <0.001 | 0.289, 0.373 |  |  |  |
| Deprivation | 0.637 | <0.001 | 0.608, 0.666 |  |  |  |
| **Outcome:** either psychopathology, executive function, or attention orienting toward angry faces |  |  |  |  |  |  |
| **Adversity:** either Threat or Deprivation |  |  |  |  |  |  |
| **Note:** Threat at baseline did not predict Deprivation at Follow-up ($\beta$=-0.036, $p$=0.115, CI95% -0.080 to 0.009), but deprivation at baseline predicted modestly lower levels of threat at follow-up (($\beta$=-0.053, $p$=0.008, CI95% -0.093 to -0.014). | | | | | | |

#### Executive Function

Executive function was moderately stable over the course of development (β=0.435, p<0.001). Cross-sectional analysis showed worse EF performance was more strongly associated with deprivation (β=-0.155, p<0.001) than threat (β=-0.086, p<0.001) at baseline and executive function was only associated with deprivation at follow-up (β=-0.061, p=0.008). Longitudinal analysis showed no significant effects of either threat or deprivation on change in executive function over the three-year follow-up (Figure 1 – Panel B).

**Figure 1.**
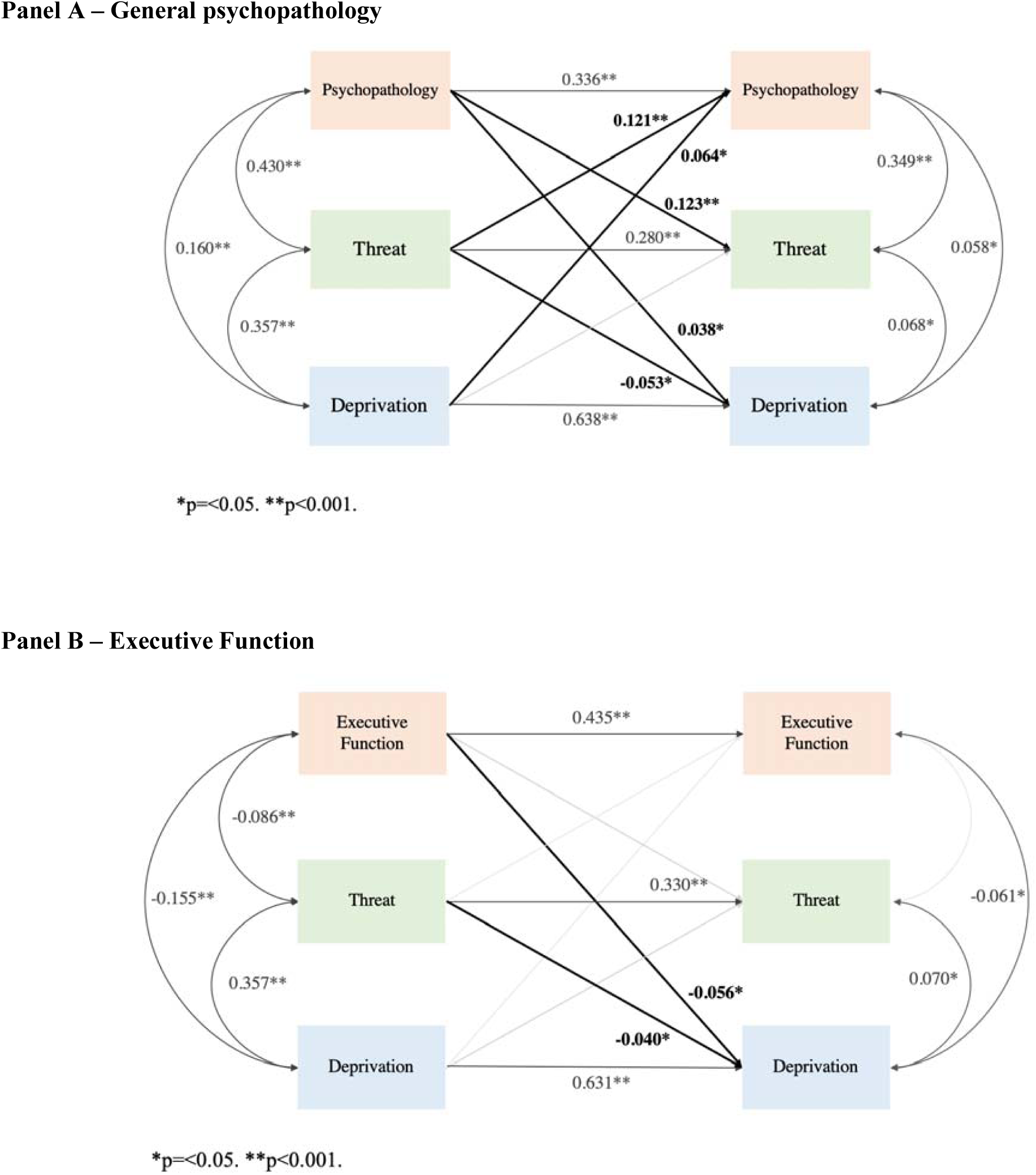
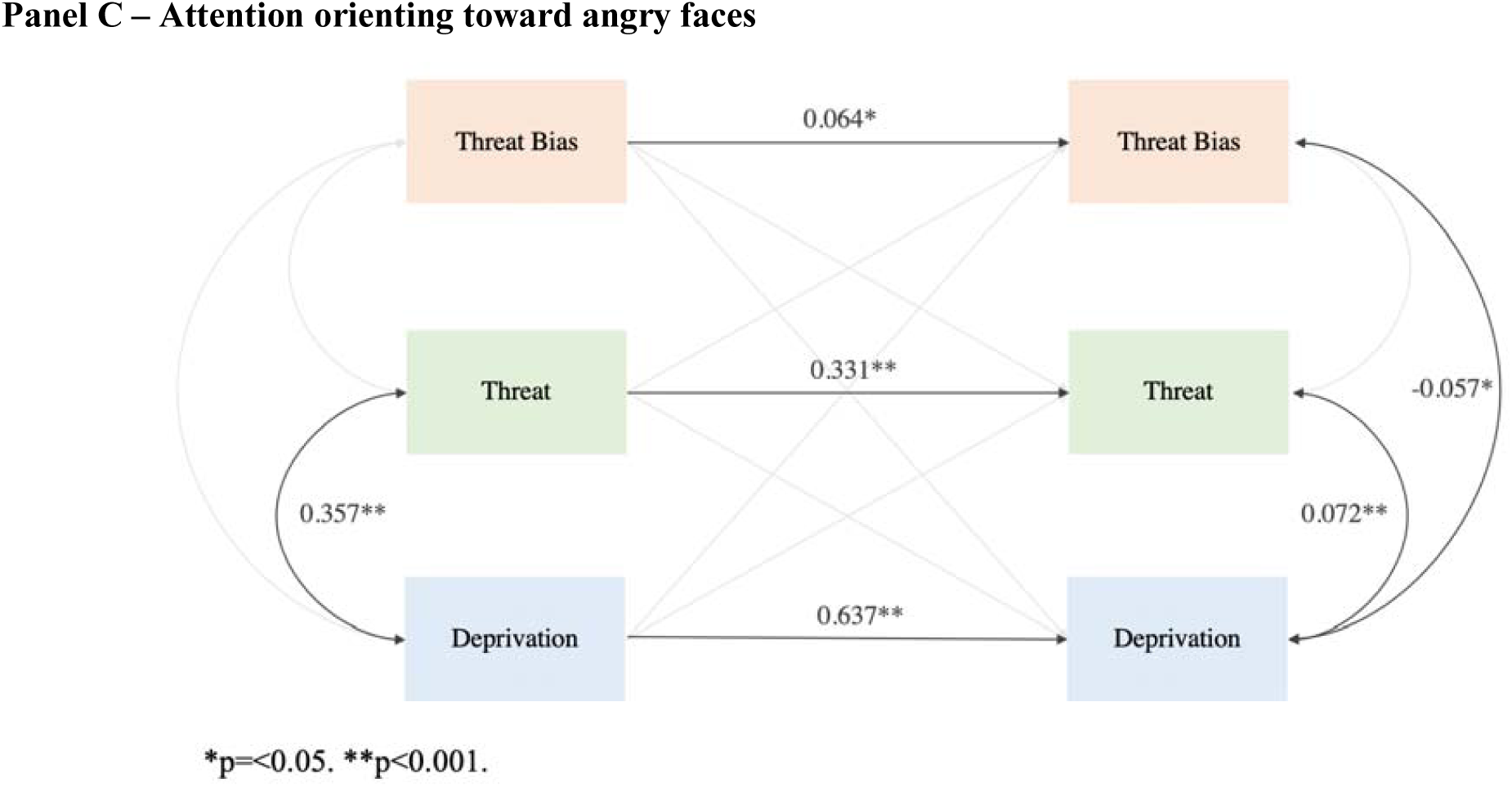
Cross Lagged Panel Models showing cross-sectional and longitudinal associations between general psychopathology, executive function, attention orienting toward angry faces and threat and deprivation in baseline and 3-year follow-up assessments

*Post hoc* analyses were conducted in order to investigate the impact of adversity on specific dimensions of EF and are shown on Table 4. All dimensions of EF were stable over the course of three years (WM, β=0.432, p<0.001; IC, β=0.310, p<0.001; TP, β=0.296, p=0.025). Cross-sectional results indicate that threat was associated with worse performance in working memory (β= −0.079, p<0.001) and temporal processing tasks (β= −0.082, p<0.001) at baseline, but not follow up. Also, deprivation was associated with all three dimensions (working memory, inhibitory control and temporal processing) of executive function at baseline (WM, β= −0.146, p<0.001; IC, β= −0.052, p<0.001; TP, β= −0.154, p=0.025), and working memory (β= −0.057, p=0.014) and inhibitory control (β= −0.067, p=0.004) at follow up. Longitudinal results indicate that deprivation, and not threat, at baseline predicted worsening performance on inhibitory control tasks over time (β= −0.086, p<0.001), but no other dimension of EF.

**Table 4.** *Post hoc* Cross-lagged Panel Models results.

| Cross-sectional |  |  |  |  |  |  |
| --- | --- | --- | --- | --- | --- | --- |
|  | Baseline |  |  | Follow-up |  |  |
| | $\beta$ | *p value* | CI 95% | $\beta$ | *p value* | CI 95% |
| Threat |  |  |  |  |  |  |
| Working memory | -0.079 | <0.001 | -0.119, -0.039 | 0.015 | 0.519 | -0.030, 0.060 |
| Inhibitory control | -0.039 | 0.057 | -0.079, 0.001 | -0.011 | 0.629 | -0.057, 0.034 |
| Temporal processing | -0.082 | <0.001 | -0.122, -0.042 | 0.019 | 0.417 | -0.027, 0.064 |
| Deprivation |  |  |  |  |  |  |
| Working memory | -0.146 | <0.001 | -0.185, -0.107 | -0.057 | 0.014 | -0.102, -0.011 |
| Inhibitory control | -0.052 | 0.010 | -0.092, -0.013 | -0.067 | 0.004 | -0.112, -0.022 |
| Temporal processing | -0.154 | <0.001 | -0.193, -0.115 | -0.044 | 0.054 | -0.090, 0.001 |
| Longitudinal associations |  |  |  |  |  |  |
|  | Adversity → Outcome |  |  | Outcome → Adversity |  |  |
| | $\beta$ | *p value* | CI 95% | $\beta$ | *p value* | CI 95% |
| Working memory |  |  |  |  |  |  |
| Threat | -0.041 | 0.064 | -0.084, 0.002 | -0.019 | 0.373 | -0.062, 0.023 |
| Deprivation | -0.023 | 0.302 | -0.067, 0.021 | -0.050 | 0.005 | -0.085, -0.015 |
| Inhibitory control |  |  |  |  |  |  |
| Threat | -0.010 | 0.659 | -0.056, 0.035 | -0.027 | 0.196 | -0.069, 0.014 |
| Deprivation | -0.086 | <0.001 | -0.132, -0.040 | -0.028 | 0.108 | -0.062, 0.006 |
| Temporal processing |  |  |  |  |  |  |
| Threat | -0.026 | 0.259 | -0.072, 0.019 | -0.033 | 0.133 | -0.076, 0.010 |
| Deprivation | -0.041 | 0.084 | -0.088, 0.006 | -0.053 | 0.004 | -0.088, -0.017 |
| Autoregressive |  |  |  |  |  |  |
|  | Autoregressive |  |  |  |  |  |
| | $\beta$ | *p value* | CC 95% | | | |
| Working memory | 0.432 | <0.001 | 0.395, 0.470 |  |  |  |
| Threat | 0.330 | <0.001 | 0.288, 0.372 |  |  |  |
| Deprivation | 0.632 | <0.001 | 0.603, 0.661 |  |  |  |
| Inhibitory control | 0.310 | <0.001 | 0.269, 0.350 |  |  |  |
| Threat | 0.330 | <0.001 | 0.288, 0.371 |  |  |  |
| Deprivation | 0.638 | <0.001 | 0.609, 0.667 |  |  |  |
| Temporal processing | 0.296 | 0.025 | 0.254, 0.339 |  |  |  |
| Threat | 0.330 | <0.001 | 0.288, 0.371 |  |  |  |
| Deprivation | 0.631 | <0.001 | 0.601, 0.660 |  |  |  |
| Outcome: either working memory, inhibitory control, and temporal processing |  |  |  |  |  |  |
| Adversity: either Threat or Deprivation |  |  |  |  |  |  |

#### Attention orienting toward angry faces

Biases toward angry faces at follow-up was significantly predicted by baseline levels, even though it was less stable than EF and general psychopathology (β= 0.064, p=0.025). Cross-sectional analysis showed deprivation was associated with bias *away* from angry faces at follow-up (β= −0.057, p=0.024,), but longitudinal analysis did not reveal any significant associations (Figure 1 – Panel C).

#### General psychopathology (the “p” factor)

General psychopathology was moderately stable over the course of three years (β=0.336, p<0.001). Cross-sectional analysis revealed psychopathology was more strongly associated with threat than with deprivation at both baseline (Threat, β=0.430, p<0.001; Deprivation, β=0.160, p<0.001) and follow-up (Threat, β=0.349, p<0.001; Deprivation, β=0.058, p=0.010). Longitudinal predictions showed that both, threat (β=0.121, p<0.001) and deprivation (β=0.064, p=0.003), predicted increases in general psychopathology three years later (Figure 1 – Panel A).

#### Bidirectional effects

Despite not being part of our primary hypothesis, our CLPM detected some bidirectional associations. We found that higher general psychopathology at baseline predicted future exposure to both threat and deprivation. Similarly, worse EF performance at baseline modestly predicted higher levels of exposure to deprivation three years later.

## Discussion

This study examined theoretical predictions of a dimensional model of childhood adversity (McLaughlin et al., 2014). Specifically, we evaluated the associations of threat and deprivation assessed dimensionally and modeled as latent factors with EF, attention orienting to threat, and psychopathology over time using a large sample in a middle-income country.

In accordance to previous findings, our results also suggest that threat and deprivation have differential associations with cognitive and emotional development, as well as psychopathology, even though not all hypotheses were supported. In particular, threat was more strongly associated with psychopathology than deprivation, and deprivation more strongly associated with EF than threat. However, threat was not uniquely associated with attention bias towards threat as expected, and instead deprivation was associated with attention bias *away* from threat. This pattern of findings has theoretical implications for conceptual models of adversity and development, as well as clinical implications regarding potential targets for early interventions aimed at preventing the long-term consequences of adversity for mental health and academic achievement.

There is mounting evidence showing that childhood adversity is associated with high levels of general psychopathology, both cross-sectionally and prospectively (Green *et al.*, 2010; Dunn *et al.*, 2017; McLaughlin et al., 2012). This link tends to span all forms of psychopathology, including both internalizing and externalizing domains (Salum *et al.*, 2016) – and therefore associations with general expressive indexes of psychopathology (such as the “p” factor) are expected (Miller *et al.*, 2016), as we confirmed here. Also, in line with some previous evidence and theoretical models (Busso, McLaughlin and Sheridan, 2017), we showed associations between adversity and psychopathology were revealed for both types of adversity domains. However, the associations of with psychopathology were stronger for threat than deprivation, which might reveal a more prominent role of the threat domain on shared expression of psychopathology, as already noted by direct and indirect effects of threat and deprivation, respectively, on psychopathology (Miller *et al.*, 2016; Platt *et al.*, 2018).

Although both threat and deprivation were each associated with worse performance on EF tasks at baseline, the associations were stronger for deprivation when compared to threat, and only deprivation predicted changes in EF over the follow-up period. This pattern is consistent with previous cross-sectional studies that observing that experiences characterized by deprivation, and not threat, are related to deficits in EF (Bos, 2009; Beckett *et al.*, 2010; Sheridan *et al.*, 2017; Machlin *et al.*, 2019; Lambert et al., 2017). In contrast to our hypotheses, neither threat nor deprivation predicted overall changes in general EF over time. However, *post hoc* analyses showed that deprivation predicted worsening performance on inhibitory control tasks over time. This suggests that different adversity types may differentially influence the development of specific domains of EF, consistent with prior work suggesting that specific types of adversity exposure might differentially impact specific cognitive abilities (Noble, McCandliss and Farah, 2007; Lambert *et al.*, 2017), and that cognitive control might be a domain of particular relevance for children who have experienced deprivation. These findings are broadly consistent with theoretical predictions arguing that deprivation may uniquely influence the development of EF in children and adolescents (McLaughlin et al., 2014; Sheridan & McLaughlin, 2014).

Different from what was expected, no associations were found between both dimensions of childhood adversity exposure and attention orienting *toward* angry faces. Conversely, a slight cross-sectional association between deprivation and attention orienting *away* from angry faces was found at follow-up. Even though previous research has already shown that children and adolescents exposed to different forms of adversity present higher attention bias toward angry faces when compared to those not exposed (Pollak and Tolley-Schell, 2003; Briggs-Gowan *et al.*, 2015; Miller, 2015), this association might depend on the developmental period of assessment. Attention bias toward threat tend to decrease from childhood through adolescence (Reinholdt-Dunne *et al.*, 2012), suggesting that with the coming of age the ability to deliberately direct attention away from threat might increase. Supporting that notion and the association between deprivation and attention bias *away* from threat found at follow up, the association between exposure to adversity and attention bias away from threat among adolescents, and not children, has already been documented (Weissman *et al.*, 2019). Such results should be interpreted with caution due to the low stability of the attention bias measure in this study, considering that the utility of using the dot-probe has already been called into question due to its low-retest reliability (Chapman, Devue and Grimshaw, 2019).

Our study has a number of strengths. First, by using a dimensional approach to childhood adversities, we were able to distinguish possible differential impacts of distinct experiences on psychopathology, EF, and attention orienting toward angry faces supporting evidence of the pathways through which adversity impacts child development. Second, our longitudinal design allowed us to explore the associations of threat and deprivation with developmental change in these domains over time, which has rarely been done in existing studies of adversity dimensions. Some limitations also should be noted. First, our study used a school-based sample assessed mainly with parent-reported events. Child-reported information would be an important piece of information to be investigated in future studies. Second, literature suggests that the deprivation dimension is characterized by cognitive stimulation, or an enriched cognitive environment (Sheridan and Mclaughlin, 2014). Our deprivation dimension did not account for any of those specific variables, suggesting that we might not be capturing deprivation as a whole. Third, attention orienting toward angry faces might captures only one relatively constrained domain of emotional processing. Because no other measure of emotional information processing was assessed in this study, we were not able to capture the effects of adversity on other domains of emotional processing argued to be particularly likely to be influenced by threat-related adversity, including emotional reactivity, emotional learning, and emotion regulation (McLaughlin & Lambert, 2017).

Exposure to adversity, especially during childhood, is a complex phenomena with meaningful and well-established effects on child development. Because adversity can take upon many forms, dimensional models—as the one investigated here—might help to disentangle the specific developmental impacts of different forms of exposure. This knowledge facilitates the identification of the risks conferred to certain individuals when exposed to certain types of adversity, as well as which interventions could be more effective in buffering these negative impacts on development. Bidirectional results suggest that reverse causality cannot be excluded, but they should be interpreted with caution given the lack of a genetically sensitive design. Therefore, further investigations on bidirectional longitudinal relationships among the variables using genetic informed models and on possible moderators and mediators for the relationships found in this paper seem to be a promising direction. Such investigations could generate more information in order to more effectively support healthy and adaptive child development through targeted interventions to high-risk populations.

## Acknowledgements

The authors thank the children and families from the Brazilian High-Risk Cohort for Psychiatric Disorders, as well as all the researchers and staff for making this study possible.

## Ethical Standards

The authors assert that all procedures contributing to this work comply with the ethical standards of the relevant national and institutional committees on human experimentation and with the Helsinki Declaration of 1975, as revised in 2008.

## Supplemental Material

### 1. Adversity

Threat and deprivation variables were selected according to documented theoretical models in order to encompass both dimensions of childhood adversity. Selected variables, their label and ranges are described below (Supplemental Table S1), as well as descriptive data for each selected variable (Supplemental Table S2). At baseline, all exposure questions were asked considering the lifetime time range, while at follow-up exposure was assessed and considered positive only if it happened over the past three-year time range.

Confirmatory factor analysis, using the full maximum likelihood to deal with missing data, were conducted using baseline and follow-up data. The threat and deprivation model at baseline was available for 2511 participants, and at follow up 2010 participants (Supplemental Figure S1).

**Supplemental Table S1.**
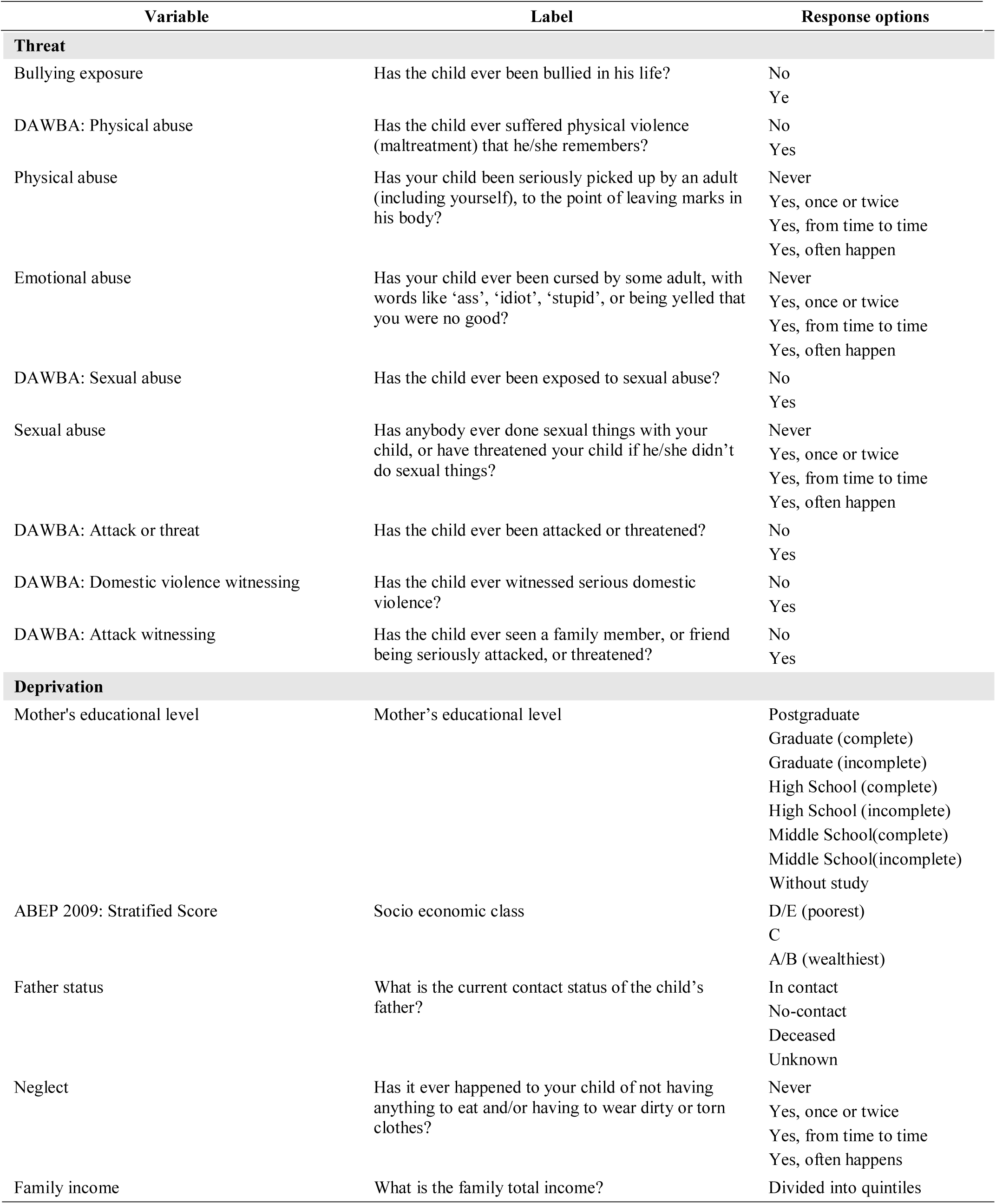
Threat and Deprivation variable description.

| Variable | Label | Response options |
| --- | --- | --- |

| Threat |  |  |
| --- | --- | --- |
| Bullying exposure | Has the child ever been bullied in his life? | No<br>Yes |
| DAWBA: Physical abuse | Has the child ever suffered physical violence (maltreatment) that he/she remembers? | No<br>Yes |
| Physical abuse | Has your child been seriously picked up by an adult (including yourself), to the point of leaving marks in his body? | Never<br>Yes, once or twice<br>Yes, from time to time<br>Yes, often happen |
| Emotional abuse | Has your child ever been cursed by some adult, with words like 'ass', 'idiot', 'stupid', or being yelled that you were no good? | Never<br>Yes, once or twice<br>Yes, from time to time<br>Yes, often happen |
| DAWBA: Sexual abuse | Has the child ever been exposed to sexual abuse? | No<br>Yes |
| Sexual abuse | Has anybody ever done sexual things with your child, or have threatened your child if he/she didn't do sexual things? | Never<br>Yes, once or twice<br>Yes, from time to time<br>Yes, often happen |
| DAWBA: Attack or threat | Has the child ever been attacked or threatened? | No<br>Yes |
| DAWBA: Domestic violence witnessing | Has the child ever witnessed serious domestic violence? | No<br>Yes |
| DAWBA: Attack witnessing | Has the child ever seen a family member, or friend being seriously attacked, or threatened? | No<br>Yes |
| Deprivation |  |  |
| Mother's educational level | Mother's educational level | Postgraduate<br>Graduate (complete)<br>Graduate (incomplete)<br>High School (complete)<br>High School (incomplete)<br>Middle School(complete)<br>Middle School(incomplete)<br>Without study |
| ABEP 2009: Stratified Score | Socio economic class | D/E (poorest)<br>C<br>A/B (wealthiest) |
| Father status | What is the current contact status of the child's father? | In contact<br>No-contact<br>Deceased<br>Unknown |
| Neglect | Has it ever happened to your child of not having anything to eat and/or having to wear dirty or torn clothes? | Never<br>Yes, once or twice<br>Yes, from time to time<br>Yes, often happens |
| Family income | What is the family total income? | Divided into quintiles |

**Supplemental Table S2.**
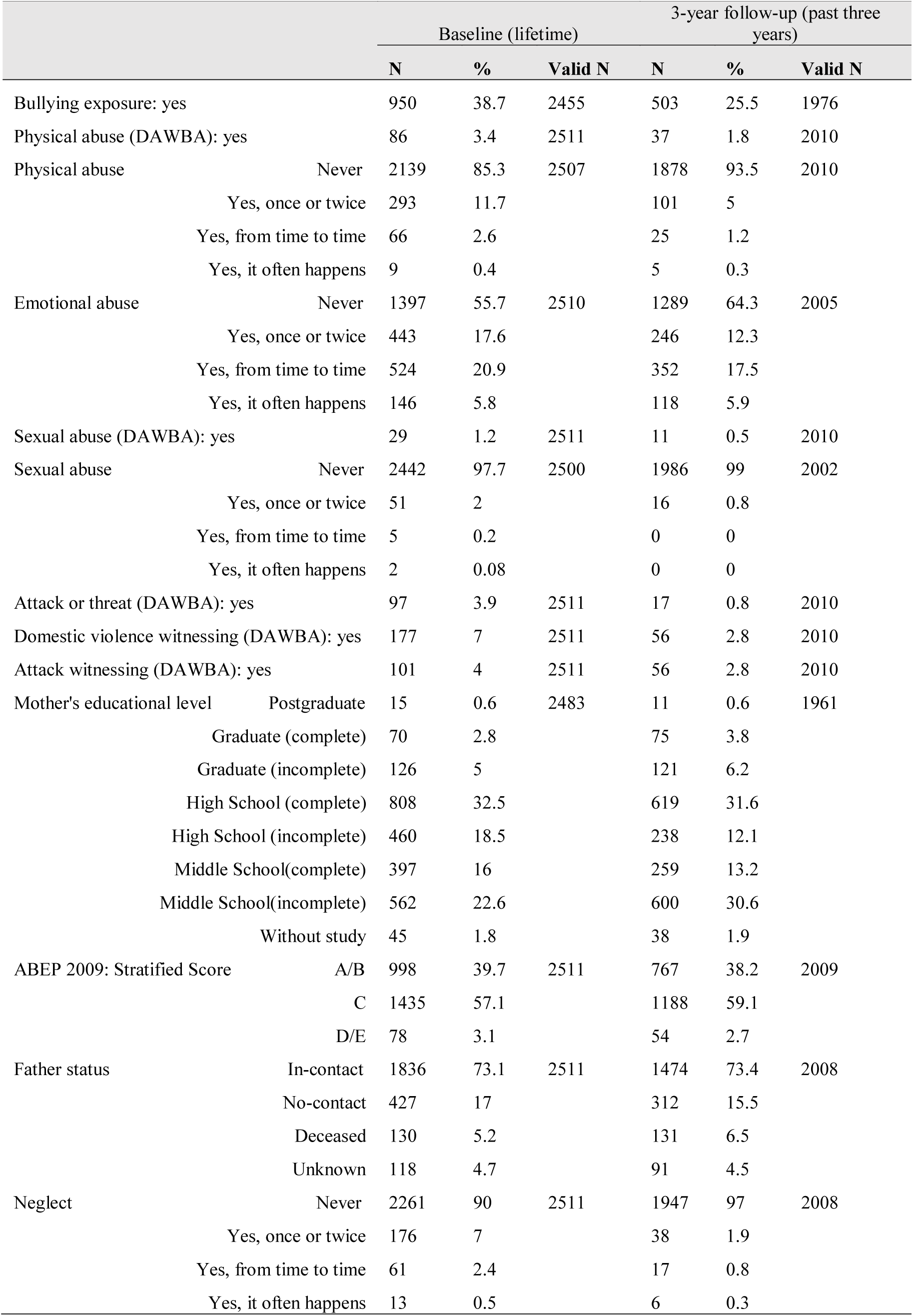
Threat and Deprivation variable frequency.

**Supplemental Figure S1.**
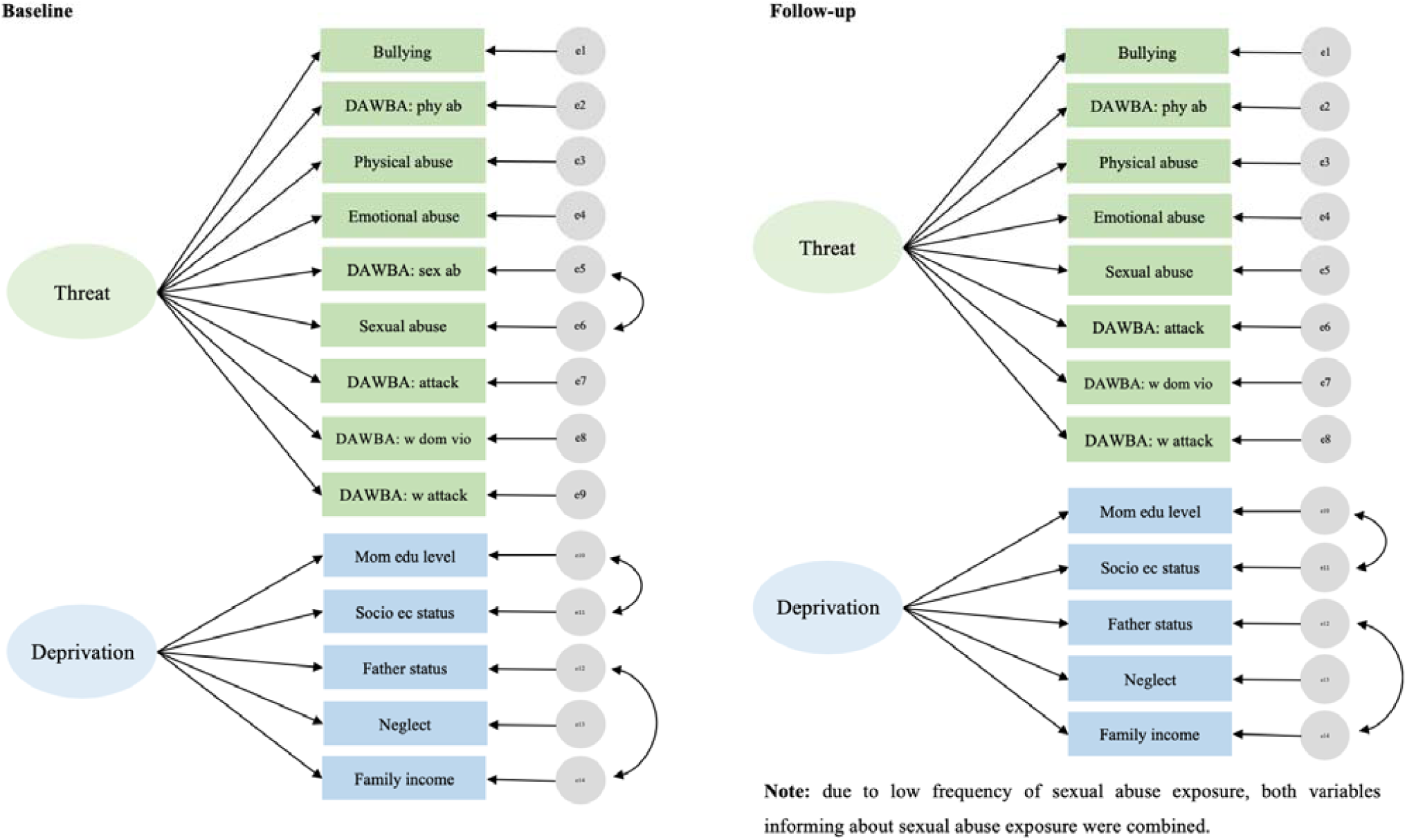
Threat and Deprivation model depiction at baseline and follow-up.

### 2. Psychopathology

Confirmatory factor analysis, using the diagonally weighted least squares (DWLS) estimator, were conducted using CBCL baseline and follow-up data using a bifactor model in which all items are loaded in a general factor (the “p” factor) and residuals variance is captured by internalizing and externalizing domains as outlined by the CBCL scoring system. The psychopathology model at baseline was available for 2511 participants and showed adequate fit indexes (CFI= 0.984, TLI= 0.983, RMSEA= 0.020). The psychopathology model at follow up was available for 2010 participants and also showed adequate fit indexes (CFI= 0.973, TLI= 0.972, RMSEA= 0.025). Factor loadings for baseline and follow-up data are found on Supplemental Table S3.

**Supplemental Table S3.**
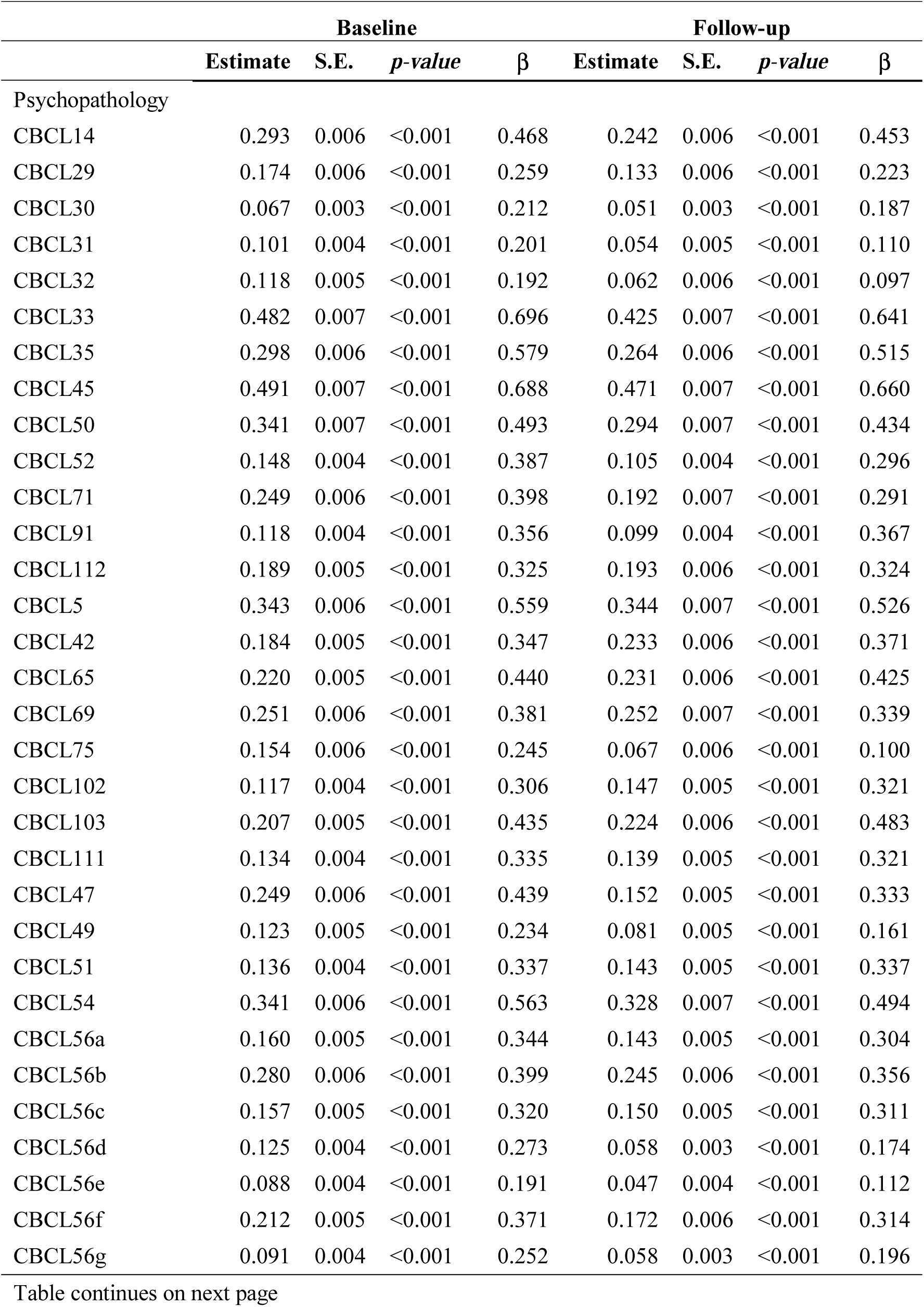
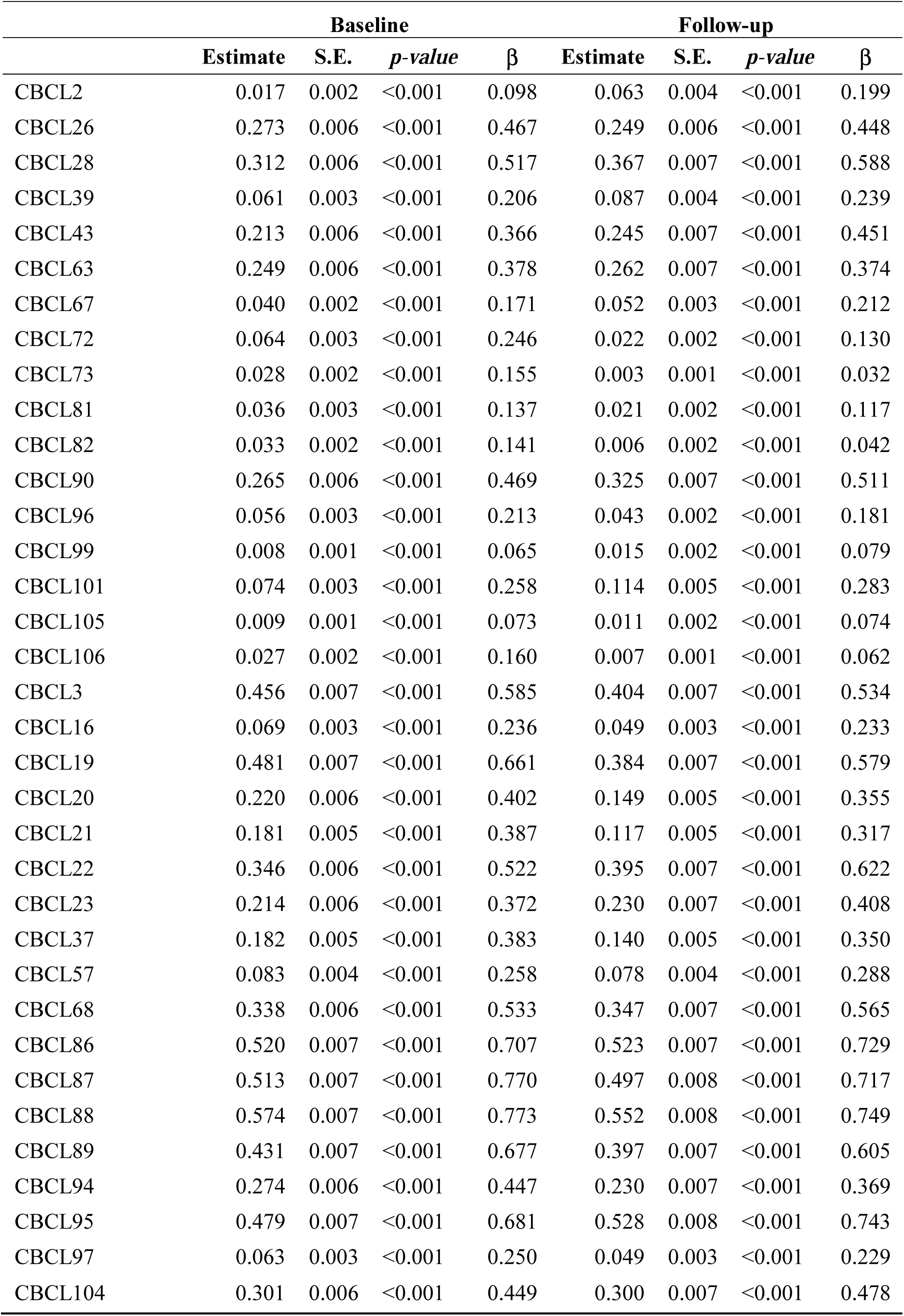
Factor loadings of the General Psychopathology Model at baseline and follow-up.

|  | Baseline |  |  |  | Follow-up |  |  |  |
| --- | --- | --- | --- | --- | --- | --- | --- | --- |
| | Estimate | S.E. | *p-value* | $\beta$ | Estimate | S.E. | *p-value* | $\beta$ |
| Psychopathology |  |  |  |  |  |  |  |  |
| CBCL14 | 0.293 | 0.006 | <0.001 | 0.468 | 0.242 | 0.006 | <0.001 | 0.453 |
| CBCL29 | 0.174 | 0.006 | <0.001 | 0.259 | 0.133 | 0.006 | <0.001 | 0.223 |
| CBCL30 | 0.067 | 0.003 | <0.001 | 0.212 | 0.051 | 0.003 | <0.001 | 0.187 |
| CBCL31 | 0.101 | 0.004 | <0.001 | 0.201 | 0.054 | 0.005 | <0.001 | 0.110 |
| CBCL32 | 0.118 | 0.005 | <0.001 | 0.192 | 0.062 | 0.006 | <0.001 | 0.097 |
| CBCL33 | 0.482 | 0.007 | <0.001 | 0.696 | 0.425 | 0.007 | <0.001 | 0.641 |
| CBCL35 | 0.298 | 0.006 | <0.001 | 0.579 | 0.264 | 0.006 | <0.001 | 0.515 |
| CBCL45 | 0.491 | 0.007 | <0.001 | 0.688 | 0.471 | 0.007 | <0.001 | 0.660 |
| CBCL50 | 0.341 | 0.007 | <0.001 | 0.493 | 0.294 | 0.007 | <0.001 | 0.434 |
| CBCL52 | 0.148 | 0.004 | <0.001 | 0.387 | 0.105 | 0.004 | <0.001 | 0.296 |
| CBCL71 | 0.249 | 0.006 | <0.001 | 0.398 | 0.192 | 0.007 | <0.001 | 0.291 |
| CBCL91 | 0.118 | 0.004 | <0.001 | 0.356 | 0.099 | 0.004 | <0.001 | 0.367 |
| CBCL112 | 0.189 | 0.005 | <0.001 | 0.325 | 0.193 | 0.006 | <0.001 | 0.324 |
| CBCL5 | 0.343 | 0.006 | <0.001 | 0.559 | 0.344 | 0.007 | <0.001 | 0.526 |
| CBCL42 | 0.184 | 0.005 | <0.001 | 0.347 | 0.233 | 0.006 | <0.001 | 0.371 |
| CBCL65 | 0.220 | 0.005 | <0.001 | 0.440 | 0.231 | 0.006 | <0.001 | 0.425 |
| CBCL69 | 0.251 | 0.006 | <0.001 | 0.381 | 0.252 | 0.007 | <0.001 | 0.339 |
| CBCL75 | 0.154 | 0.006 | <0.001 | 0.245 | 0.067 | 0.006 | <0.001 | 0.100 |
| CBCL102 | 0.117 | 0.004 | <0.001 | 0.306 | 0.147 | 0.005 | <0.001 | 0.321 |
| CBCL103 | 0.207 | 0.005 | <0.001 | 0.435 | 0.224 | 0.006 | <0.001 | 0.483 |
| CBCL111 | 0.134 | 0.004 | <0.001 | 0.335 | 0.139 | 0.005 | <0.001 | 0.321 |
| CBCL47 | 0.249 | 0.006 | <0.001 | 0.439 | 0.152 | 0.005 | <0.001 | 0.333 |
| CBCL49 | 0.123 | 0.005 | <0.001 | 0.234 | 0.081 | 0.005 | <0.001 | 0.161 |
| CBCL51 | 0.136 | 0.004 | <0.001 | 0.337 | 0.143 | 0.005 | <0.001 | 0.337 |
| CBCL54 | 0.341 | 0.006 | <0.001 | 0.563 | 0.328 | 0.007 | <0.001 | 0.494 |
| CBCL56a | 0.160 | 0.005 | <0.001 | 0.344 | 0.143 | 0.005 | <0.001 | 0.304 |
| CBCL56b | 0.280 | 0.006 | <0.001 | 0.399 | 0.245 | 0.006 | <0.001 | 0.356 |
| CBCL56c | 0.157 | 0.005 | <0.001 | 0.320 | 0.150 | 0.005 | <0.001 | 0.311 |
| CBCL56d | 0.125 | 0.004 | <0.001 | 0.273 | 0.058 | 0.003 | <0.001 | 0.174 |
| CBCL56e | 0.088 | 0.004 | <0.001 | 0.191 | 0.047 | 0.004 | <0.001 | 0.112 |
| CBCL56f | 0.212 | 0.005 | <0.001 | 0.371 | 0.172 | 0.006 | <0.001 | 0.314 |
| CBCL56g | 0.091 | 0.004 | <0.001 | 0.252 | 0.058 | 0.003 | <0.001 | 0.196 |
| | Estimate | S.E. | *p-value* | $\beta$ | Estimate | S.E. | *p-value* | $\beta$ |
| CBCL2 | 0.017 | 0.002 | <0.001 | 0.098 | 0.063 | 0.004 | <0.001 | 0.199 |
| CBCL26 | 0.273 | 0.006 | <0.001 | 0.467 | 0.249 | 0.006 | <0.001 | 0.448 |
| CBCL28 | 0.312 | 0.006 | <0.001 | 0.517 | 0.367 | 0.007 | <0.001 | 0.588 |
| CBCL39 | 0.061 | 0.003 | <0.001 | 0.206 | 0.087 | 0.004 | <0.001 | 0.239 |
| CBCL43 | 0.213 | 0.006 | <0.001 | 0.366 | 0.245 | 0.007 | <0.001 | 0.451 |
| CBCL63 | 0.249 | 0.006 | <0.001 | 0.378 | 0.262 | 0.007 | <0.001 | 0.374 |
| CBCL67 | 0.040 | 0.002 | <0.001 | 0.171 | 0.052 | 0.003 | <0.001 | 0.212 |
| CBCL72 | 0.064 | 0.003 | <0.001 | 0.246 | 0.022 | 0.002 | <0.001 | 0.130 |
| CBCL73 | 0.028 | 0.002 | <0.001 | 0.155 | 0.003 | 0.001 | <0.001 | 0.032 |
| CBCL81 | 0.036 | 0.003 | <0.001 | 0.137 | 0.021 | 0.002 | <0.001 | 0.117 |
| CBCL82 | 0.033 | 0.002 | <0.001 | 0.141 | 0.006 | 0.002 | <0.001 | 0.042 |
| CBCL90 | 0.265 | 0.006 | <0.001 | 0.469 | 0.325 | 0.007 | <0.001 | 0.511 |
| CBCL96 | 0.056 | 0.003 | <0.001 | 0.213 | 0.043 | 0.002 | <0.001 | 0.181 |
| CBCL99 | 0.008 | 0.001 | <0.001 | 0.065 | 0.015 | 0.002 | <0.001 | 0.079 |
| CBCL101 | 0.074 | 0.003 | <0.001 | 0.258 | 0.114 | 0.005 | <0.001 | 0.283 |
| CBCL105 | 0.009 | 0.001 | <0.001 | 0.073 | 0.011 | 0.002 | <0.001 | 0.074 |
| CBCL106 | 0.027 | 0.002 | <0.001 | 0.160 | 0.007 | 0.001 | <0.001 | 0.062 |
| CBCL3 | 0.456 | 0.007 | <0.001 | 0.585 | 0.404 | 0.007 | <0.001 | 0.534 |
| CBCL16 | 0.069 | 0.003 | <0.001 | 0.236 | 0.049 | 0.003 | <0.001 | 0.233 |
| CBCL19 | 0.481 | 0.007 | <0.001 | 0.661 | 0.384 | 0.007 | <0.001 | 0.579 |
| CBCL20 | 0.220 | 0.006 | <0.001 | 0.402 | 0.149 | 0.005 | <0.001 | 0.355 |
| CBCL21 | 0.181 | 0.005 | <0.001 | 0.387 | 0.117 | 0.005 | <0.001 | 0.317 |
| CBCL22 | 0.346 | 0.006 | <0.001 | 0.522 | 0.395 | 0.007 | <0.001 | 0.622 |
| CBCL23 | 0.214 | 0.006 | <0.001 | 0.372 | 0.230 | 0.007 | <0.001 | 0.408 |
| CBCL37 | 0.182 | 0.005 | <0.001 | 0.383 | 0.140 | 0.005 | <0.001 | 0.350 |
| CBCL57 | 0.083 | 0.004 | <0.001 | 0.258 | 0.078 | 0.004 | <0.001 | 0.288 |
| CBCL68 | 0.338 | 0.006 | <0.001 | 0.533 | 0.347 | 0.007 | <0.001 | 0.565 |
| CBCL86 | 0.520 | 0.007 | <0.001 | 0.707 | 0.523 | 0.007 | <0.001 | 0.729 |
| CBCL87 | 0.513 | 0.007 | <0.001 | 0.770 | 0.497 | 0.008 | <0.001 | 0.717 |
| CBCL88 | 0.574 | 0.007 | <0.001 | 0.773 | 0.552 | 0.008 | <0.001 | 0.749 |
| CBCL89 | 0.431 | 0.007 | <0.001 | 0.677 | 0.397 | 0.007 | <0.001 | 0.605 |
| CBCL94 | 0.274 | 0.006 | <0.001 | 0.447 | 0.230 | 0.007 | <0.001 | 0.369 |
| CBCL95 | 0.479 | 0.007 | <0.001 | 0.681 | 0.528 | 0.008 | <0.001 | 0.743 |
| CBCL97 | 0.063 | 0.003 | <0.001 | 0.250 | 0.049 | 0.003 | <0.001 | 0.229 |
| CBCL104 | 0.301 | 0.006 | <0.001 | 0.449 | 0.300 | 0.007 | <0.001 | 0.478 |

### 3. Executive Function

Six executive function tasks were used as measures of working memory, inhibitory control and temporal processing. Missing data differs from one task to another over both time points due to assessment being performed over four sessions at baseline.

Executive function was derived from a second order model encompassing working memory inhibitory control and temporal processing at baseline and follow up. Due to the differing variability of the executive function tasks, scores were standardized regressing using General Additive Models regressing out effects of age and gender on the task parameters. At baseline, each executive function dimension was encompassed by two cognitive tasks, and at follow-up, due to the absence of one of the tasks data, the temporal processing dimension was encompassed by one task (Supplemental Figure S2).

Confirmatory factor analysis, using the full maximum likelihood to deal with missing data, were conducted using baseline and follow-up data. The executive function model at baseline was available for 2396 participants, and the follow up model for 1880 participants. Fit indexes and factor loadings are shown on Supplemental Table S4.

**Supplemental Table S4.**
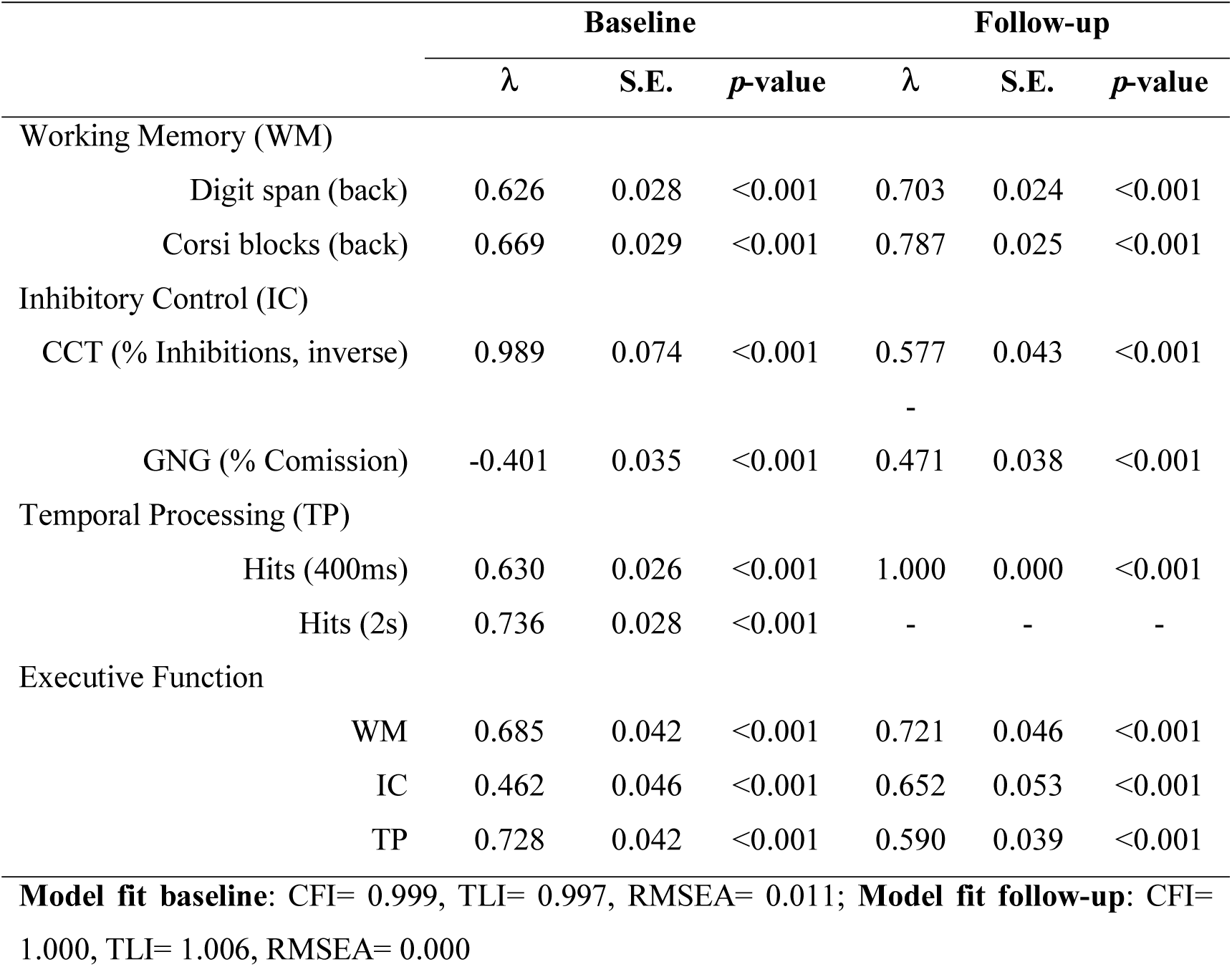
Standardized factor loadings of the baseline Executive Function Model.

**Supplemental Figure S2.**
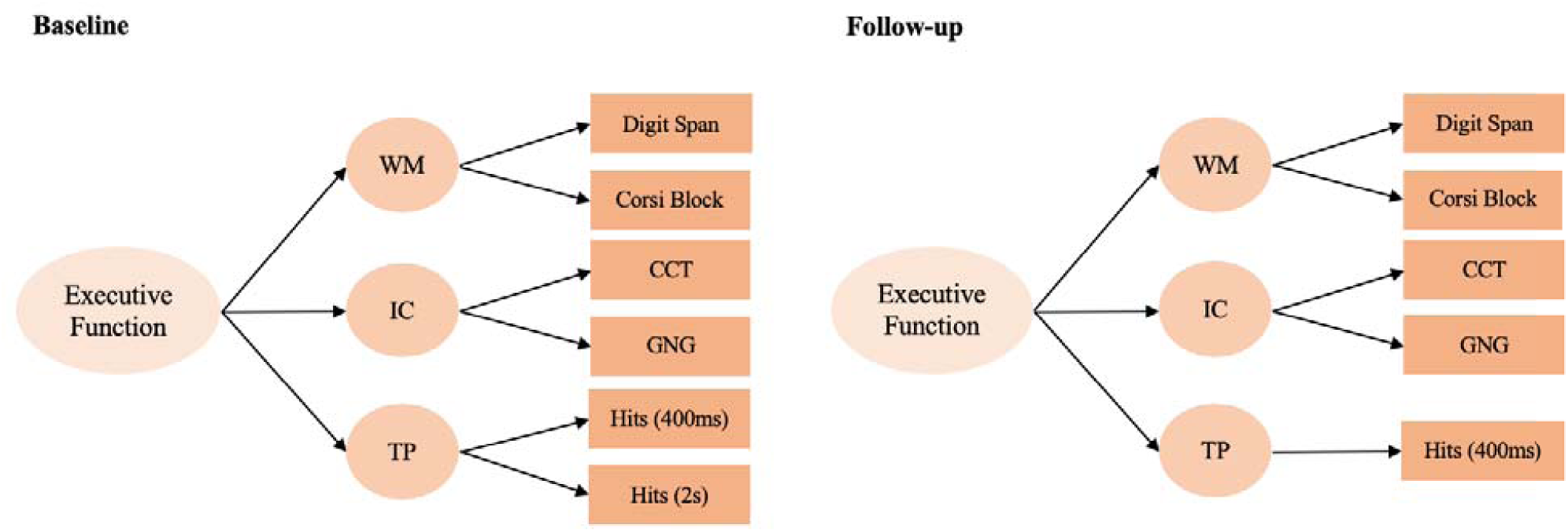
Executive Function Model baseline and follow=up.

